# Fate of glair glands and spawning behaviour in unmated crayfish: a comparison between gonochoristic slough crayfish and parthenogenetic marbled crayfish

**DOI:** 10.1101/047654

**Authors:** Günter Vogt

**Affiliations:** Faculty of Biosciences, University of Heidelberg, Im Neuenheimer Feld 230, 69120 Heidelberg, Germany

**Keywords:** freshwater crayfish, glair glands, spawning, mating, *Procambarus fallax*, *Procambarus virginalis*

## Abstract

In the period before spawning, freshwater crayfish females develop glair glands on the underside of the pleon. These glands produce the mucus for a transient tent-like structure in which the eggs are fertilized and attached to the pleopods. Long-term observation of females of the bisexually reproducing slough crayfish, *Procambarus fallax*, kept in captivity revealed that glair glands developed in late winter and late summer of each year independent of the presence of males. However, in contrast to mated females unmated females did neither form a fertilization tent nor spawn. Their glair glands persisted for an unusually long period of time and disappeared only during the next moult. Apparently, females use information on sperm availability to either reproduce or save the resources. Marbled crayfish, *Procambarus virginalis*, a parthenogenetic descendant of slough crayfish, developed glair glands in approximately the same periods of the year but spawned despite of the absence of mating. These findings indicate that on their way from gonochorism to parthenogenesis regulation of glair gland activity and spawning has been decoupled from mating. Therefore, the species pair *Procambarus fallax/Procambarus virginalis* seems to be particularly suitable to investigate the physiological, molecular and genetic mechanisms underlying spawning in freshwater crayfish.

## Introduction

In the weeks before spawning, freshwater crayfish develop prominent whitish glair glands on the underside of the pleon (Reynolds 2002). These glands first appear as faint creamy-whitish patches and become more and more distinct, being good indicators of forthcoming spawning (Jussila et al. 1996). Shortly before egg laying the glair glands release translucent mucus that forms a tent like structure on the underside of the thorax and pleon (Andrews 1904; Vogt 2016). Within this gelatinous mass the sperm is mobilised from the spermatophores that are either externally attached or internally stored, depending on crayfish family (Reynolds 2002; Vogt 2002; Niksirat et al. 2013). The fertilization tent further secures attachment of the fertilized eggs to the pleopods, where they are brooded for weeks until hatching of the first juvenile stage and beyond (Vogt and Tolley 2004). Glair glands and fertilization tent are unique features of freshwater crayfish and occur in all three families, the Astacidae, Cambaridae and Parastacidae (Andrews 1906; Mason 1970; Thomas and Crawley 1975; Jussila et al. 1996; Vogt 2016). Simpler cement glands that only facilitate attachment of the eggs to the pleopods also occur in other Decapoda (Adiyodi and Anilkumar 1988).

During crustacean conferences and student courses I was often confronted with the question what happens with the glair glands and the mature oocytes if crayfish females remain unmated. Unfortunately, there is no published data on the fate of the glair glands and only little data on spawning in unmated females (Woodlock and Reynolds 1988; Reynolds et al. 1992; Holdich et al. 1995; Tropea et al. 2010; Buřič et al. 2011). One possibility is that unmated females spawn like mated females but the unfertilized eggs decay. Alternatively, the mucus and mature oocytes may be reabsorbed to avoid wasting of resources. In the present communication, I report on the fate of the glair glands and spawning behaviour in laboratory-raised mated and unmated slough crayfish *Procambarus fallax*, and their parthenogenetically reproducing relative, the marbled crayfish *Procambarus virginalis*. Marbled crayfish have no males and originated from slough crayfish by autotriploidization and concomitant transition from gonochorism to parthenogenesis (Scholtz et al. 2003; Martin et al. 2010; 2016; Vogt et al. 2015). They were recently recognized as a separate species due to reproductive and geographic isolation from *Procambarus fallax*, unique genetic and epigenetic signatures and superior growth and fecundity (Vogt et al. 2015).

## Materials and methods

Females of slough crayfish *Procambarus fallax* (Hagen 1870) (Decapoda: Cambaridae) were separated from their male batch-mates in juvenile stage 5, the first life stage that enables external sex determination. In the following five months they were kept communally in a female group and thereafter individually. Three of these females remained unmated and another three females were mated with males during the reproduction seasons in spring 2015, autumn 2015 and spring 2016. Mating was done by pairing individual females with glair glands and individual males with hooks on the ischia of the 3rd and 4th pereiopods (indicators of sexual maturity) in a 30 × 25 × 20 cm container. Copulation was considered successful when the males had turned the females on the back and inserted their copulatory organs into the sperm receptacle (annulus ventralis) of the female for at least 30 minutes. All mated and unmated females were kept individually in plastic containers of 30 × 25 × 20 cm equipped with gravel and shelter. Tap water was used as the water source and replaced once a week. Water temperature fluctuated from ∼15°C in winter to ∼25°C in summer and photoperiod was the natural ambient regimen of Heidelberg (Germany). The crayfish were fed daily ad libitum with TetraWafer Mix pellets. The observation period lasted from July 1, 2014, to May 31, 2016.

The time periods in which glair glands were visible were recorded. I have given the time in weeks only because at the beginning of glair gland maturation it is difficult to decide whether development has already started or not. The data on *Procambarus fallax* were compared with my records on the presence of glair glands in the parthenogenetic marbled crayfish, *Procambarus virginalis*, which I have collected over the last 13 years. Marbled crayfish were raised under the same conditions as slough crayfish, either individually or communally.

## Results

The females of both crayfish species developed glair glands in the reproduction seasons. These glands were located in the last thoracic sterna, the sterna and pleura of the pleon, the uropods and the pleopods (Figure 1A, B). The glair glands developed autonomously in the absence of males.

**Figure 1.**
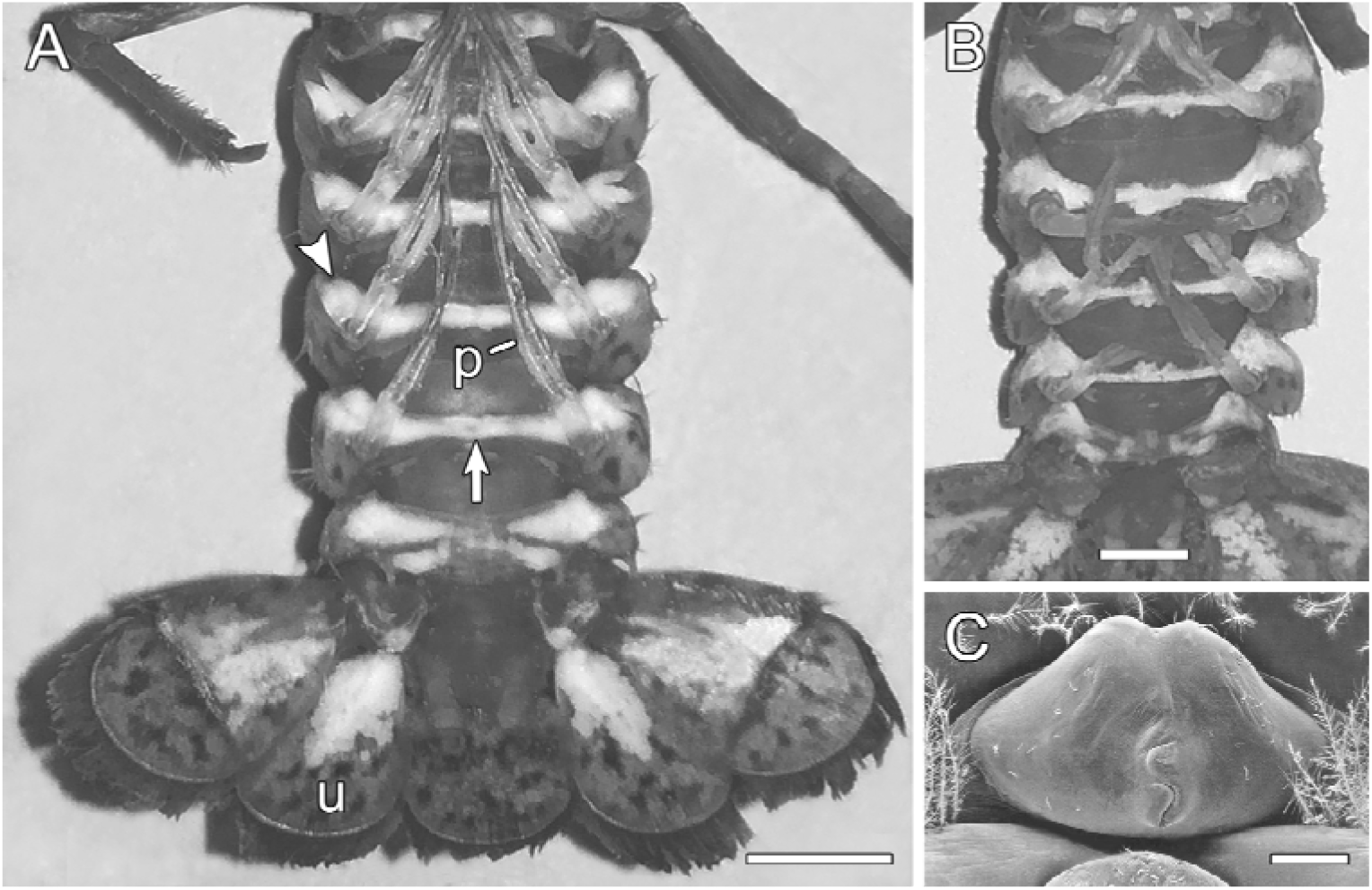
Glair glands and sperm receptacle in crayfish. (A) Mature glair glands on underside of pleon of parthenogenetic *Procambarus virginalis* female. The whitish glands are located in the sterna (arrow), pleura (arrowhead), uropods (u) and pleopods (p). Bar: 5 mm. (B) Pleon of unmated *Procambarus fallax* female showing glair glands 12 weeks after their first appearance. Bar: 2 mm. (C) Scanning electron micrograph of annulus ventralis of *Procambarus virginalis* female. Bar: 200 µm.

In the three reproduction periods analysed, unmated females of *Procambarus fallax* did not spawn although they regularly developed glair glands (Table 1). In contrast, the mated females generally laid eggs. In the latter, the glair glands were visible for about three to five weeks. They were emptied shortly before egg-laying to form the fertilization tent. In unmated females the glair glands persisted up to 12 weeks (Table 1) and disappeared only during the next ecdysis.

**Table 1.** Fate of glair glands and spawning behaviour in unmated and mated *Procambarus fallax* females.

| Females | Spring 2015 |  | Autumn 2015 |  | Spring 2016 |  |
| --- | --- | --- | --- | --- | --- | --- |
|  | glair glands | spawned | glair glands | spawned | glair glands | spawned |
| Unmated (n=3) | 7-8 wk | — | 6-9 wk | — | 7-12 wk | — |
| Mated (n=3) | 3-4 wk | + | 3-4 wk | + | 3-5 wk | + |

Laboratory reared marbled crayfish mostly reproduced twice a year like slough crayfish (Vogt 2015). Some individuals reproduced only once or three times per year. They showed glair glands with increasing staining intensity 3-6 weeks prior to spawning. In contrast to unmated slough crayfish females, parthenogenetic marbled crayfish females produced fertilization tents and spawned. Only in one out of 103 documented cases a female with glair glands skipped spawning and moulted instead. The glair glands of this specimen disappeared during moulting as in unmated slough crayfish.

## Discussion

Comparison of unmated females of bisexually reproducing slough crayfish and parthenogenetic marbled crayfish revealed interesting differences with respect to spawning and the fate of the glair glands. Mated slough crayfish females and marbled crayfish females usually spawn twice a year (Vogt 2015). These spawnings are associated with emptying of the glair glands. In slough crayfish both events are dependent on mating. Unmated *Procambarus fallax* females neither spawn nor empty their glair glands. Instead, the glair glands persist for many more weeks, probably in expectation of a later mating chance. This is in contrast to marbled crayfish which develop glair glands and spawn in the absence of males. Obviously, the information system that allows distinction between mated and unmated conditions has been lost or changed during transition from gonochorism to parthenogenesis in the *Procambarus fallax*/*Procambarus virginalis* species pair.

Complete inhibition of spawning was also observed in unmated females of the astacids *Austropotamobius pallipes*, *Astacus leptodactylus* and *Pacifastacus leniusculus* by Holdich et al. (1995). In contrast, Woodlock and Reynolds (1988) observed normal spawning but decay of the attached eggs in unmated females of *Austropotamobius pallipes* that were held in female pairs. In unmated females of laboratory-raised *Cherax destructor* (Parastacidae) the spawning behaviour was quite diverse: about 20% spawned when raised at 30°C but none at 27°C (Tropea et al. 2010). Spawning of unmated females also occurred in facultatively parthenogenetic females of the cambarid *Orconectes limosus* (Buřič et al. 2011, 2013). Out of 30 unmated parthenogenetic females 28 spawned and only one female did not spawn (the remaining individual died) (Buřič et al. 2011), resembling the situation in marbled crayfish. The reasons for the differences in spawning behaviour of unmated females between and within the cited sexually reproducing species are unknown but may rely on species differences, environmental conditions, pre-experimental experiences and the nutritional status of the individuals. Information on the fate of the glair glands is lacking in all of the above cited reports. The inhibition of glair secretion and spawning may be interpreted as an energy saving strategy.

Maturation of the oocytes is independent from the presence of males as shown by all of the above cited studies. The same holds for maturation of the glair glands which develop in the weeks before spawning. Both processes are apparently closely linked and elicited by environmental cues, mainly temperature and photoperiod (Reynolds 2002; Vogt 2015), which influence the hormonal system. Ovarian maturation is inhibited by the vitellogenesis-inhibiting hormone (VIH) from the X-organ-sinus gland system and promoted by the ovary-stimulating hormone (OSH) from the thoracic nerve ganglia (Reynolds 2002; Vogt 2002). Maturation of the glair glands is probably regulated by the same hormones.

Mating may be one of the triggers for spawning as assumed by Reynolds (2002). This assumption is supported by the experiments of Holdich et al. (1995) who paired females with males from another species. Such females generally spawned but the eggs did not develop. However, under natural conditions there are often many days or even weeks between mating and spawning suggesting that spawning could also depend on the remembrance of mating by the female or sensing of the presence of sperm. In astacid and parastacid crayfish, the spermatophores are externally attached to the thoracic sternal plates and the tail fan (Reynolds 2002; Vogt 2002; Niksirat et al. 2013), and thus, females could easily sense the presence of sperm with their pereiopods. In Cambaridae, sperm is stored internally in the annulus ventralis (Figure 1C) (Reynolds 2002; Vogt et al. 2004), and in this case, mechanoreceptors might give information on its filling status.

It is unknown whether crayfish can voluntarily empty the glair glands and ovaries by contraction of the glandular and ovarian musculatures. It is more likely, that both processes are involuntary and regulated by hormones like prostaglandin F_2α_, which was shown to increase sharply in the final stage of oogenesis in *Procambarus paeninsulanus* (Spaziani et al. 1995). In vitro tests revealed that this hormone elicits contraction of the ovarian tissue in a dose dependent manner.

The present study is the first report on the fate of the glair glands in unmated crayfish raised under controlled conditions. The differences between gonochoristic *Procambarus fallax* and parthenogenetic *Procambarus virginalis* makes this crayfish pair an interesting model system for researchers interested in the physiology, molecular biology and genetics of spawning in crayfish. The availability of a complete and searchable draft genome of the marbled crayfish (http://marmorkrebs.dkfz.de/) and complete genomes of *Procambarus fallax* females and males should facilitate research in this direction tremendously. An interesting complementary model is *Orconectes limosus* which can reproduce by gonochorism or parthenogenesis depending on conditions. In the *Procambarus fallax/Procambarus virginalis* pair the shift between gonochorism and parthenogenesis is irreversible and thus the differences in spawning should be genetically determined whereas the facultative shift between gonochorism and parthenogenesis in *Orconectes limosus* should be epigenetically determined.

## Acknowledgements

Many thanks to Ute Bieberstein for taking Figures 1A and B.

